# Whole-genome polymorphisms and relatedness of rice varieties circulating in the Mediterranean market

**DOI:** 10.1101/2025.05.29.656782

**Authors:** Hugo M. Rodrigues, M. Beatriz Vieira, Pedro M. Barros, M. Margarida Oliveira

**Author notes:** equal contribution (Hugo M. Rodrigues, M. Beatriz Vieira).

## Abstract

Rice cultivation in Europe is declining as consumers increasingly prefer imported exotic varieties, such as aromatic and basmati rice, which are prone to fraudulent varietal claims due to their higher market value. To address this issue, we sequenced 20 high-value rice cultivars circulating in the Mediterranean market, analyzing their phylogeny and whole-genome polymorphisms. Our results revealed that two basmati varieties are genetically closer to two Mediterranean varieties, and that no direct link exists between genome-wide single nucleotide polymorphism (SNP) patterns and the rice commercial category. We further discuss genes located in previously described quantitative trait loci (QTLs) related to eating quality and seed properties. A variant in the *WX1* gene, associated with higher amylose content, was found in both the Basmati group and Mediterranean varieties. Additionally, a SNP that could disrupt the drought tolerance gene *OsbHLH148* was identified in five European varieties, while a variant affecting splicing of *OsPol lambda*, related to drought response, was present in four of those. This data could assist certification offices in reducing fraud in the rice market and provide valuable insights for researchers and breeders, particularly regarding the production, consumption, and adaptation of these cultivars to the Mediterranean region.

## Introduction

As the primary staple food for over half of the world’s population, rice (*Oryza sativa*) accounts for about 20% of the calories consumed worldwide ^1^. Originating in Asia, *Oryza sativa* is highly adaptable to different latitudes and longitudes and is currently grown worldwide in a vast range of ecosystems. Genetic structure studies have revealed two primary subgroups within the *O. sativa* species: indica and japonica, arising from independent domestication events, but other types with less clear origins as sadri-basmati and aus-boro have also been considered ^2,3^.

Rice cultivation in Europe has a relatively short history, and a slight decline in rice production has been reported in recent years across this continent ^4^. While the European pedigree mainly consists of temperate japonica varieties, European consumer preferences tendentially lean towards characteristics related to grain size, colour, and cooking qualities, with a growing demand for aromatic and indica varieties ^4–6^. This, coupled with increasing prices for japonica rice, has led to a rise in imports of “exotic” varieties, and the EU has become a net importer of rice, with about one-third of its consumption sourced from countries like Pakistan, India, and Thailand ^6^. Consequently, efforts are underway to boost EU japonica rice production and sales within the European market, targeting varieties specifically produced and grown in the European region. Furthermore, the influx of new/exotic accessions is fostering fraudulent varietal claims of higher quality and expensive rice (as the example of basmati rice), with replacement or mixture of lower quality and cheaper varieties ^7,8^.

The challenges imposed by climate change and the increasing human population also drive the necessity for better, more resilient, and more nutritious rice varieties. According to “The Second Report on the State of the World’s Plant Genetic Resources for Food and Agriculture”, there are over 700,000 certified varieties distributed in genebanks worldwide ^9^. The 3K Rice Genome Project ^10^ was launched to better understand the genetic diversity of this broad germplasm, by obtaining the whole-genome sequence of more than 3000 target accessions. The data produced in this project serves as an unparalleled resource for uncovering rice genetic variation on a large scale ^1^. From this initiative, 29 million single nucleotide polymorphisms (SNPs), 2.4 million small insertions and deletions (InDels), and over 90,000 structural variations (SVs) were identified ^1^. The public availability of such genomic data allows for increased knowledge of rice populations, varieties, and even other species from the *Oryza* genus. Second ^11^ discussed the application of molecular markers to perform phylogenetic analysis in rice and, in 1999, Ge et al. performed a phylogenetic analysis from well-described genes, using only 2 nuclear genes (*Adh1* and *Adh2*) and 1 from the chloroplast (*matK*). Since then, the evolution of technology helped to detect and target the smallest genomic variations and use them to either analyse population structure or possibly implement fraud detection methods ^7,13,14^. Moreover, molecular markers have facilitated breeding programs targeting not only agronomic but also eating and cooking quality traits, and have been widely applied for genomic studies. One example is the multiple polymorphisms associated with the *waxy* gene (acting in amylose synthesis), which have been applied to assess the authenticity of carnaroli rice and estimate amylose levels in different rice varieties ^15–17^.

Our study aimed to characterize the genetic background of 22 rice varieties currently circulating in the Mediterranean market, and generate knowledge to further tackle fraudulent varietal claims and contribute to genotype conservation. The data generated highlighted the genetic relatedness of these varieties and genetic variability within genes of interest regarding grain-related traits. These data may also be used for the design of reliable and cost-effective DNA-based adulteration detection methods.

## Materials and Methods

### Plant material and DNA extraction

This study targeted a total of 22 Mediterranean rice varieties (Table 1). Two of them were previously studied by Reig-Valiente^14^ and their genome sequences retrieved from ENA (PRJEB13328): Bomba, under the accession replicates SAMEA3927584, SAMEA3927585, and SAMEA3927584; and Puntal, with accession replicates SAMEA3927614, SAMEA3927615, and SAMEA3927616. For each of the additional 20 varieties, rice seeds were germinated in hydroponics for a period of 10-16 days. The shoots of about 25 seedlings (two weeks old) were harvested and immediately frozen in liquid nitrogen for storage at -80 ºC. Frozen shoots were ground in liquid nitrogen and used for DNA extraction using an optimized CTAB-based method ^18^ with an increase of RNAse final concentration to 20 ug/mL. DNA from the Basmati Type III variety was exceptionally extracted from seed flour following CTAB extraction as described in ^19^, after the seeds were dehusked and ground into a fine powder, using a disinfected coffee grinder.

**Table 1.** List of varieties selected from those produced in the Mediterranean region and/or circulating in the European market, with respective commercial type, country of origin, and relevance in terms of country/region with higher production/consumption of the variety

| Commercial type | Variety | Origin | Relevant in |
| --- | --- | --- | --- |
| Short grain | * Bomba | Spain | Spain |
|  | Gageron | France | France |
|  | Giza 177 | Egypt | Egypt |
| Medium grain | JSendra | Spain | Spain |
|  | Manobi | France | France |
| Long A | Albatros | Italy | Portugal |
|  | Arborio | Italy | Italy |
|  | Arelate | France | France |
|  | Ariete | Italy | Portugal |
|  | Caravela | Portugal | New variety (registered in 2021) |
|  | Carnaroli | Italy | Italy |
|  | Giza 181 | Egypt | Egypt |
|  | Lusitano | Italy | Portugal |
|  | Ronaldo | Italy | Portugal |
|  | Teti | Italy | Portugal |
|  | Ulisse | Italy | Italy |
| Long B | CL-28 | Italy | Portugal |
|  | Maçarico | Portugal | Portugal |
|  | * Puntal | Spain | Spain |
| European Aromatic | Elettra | Italy | Italy |
| Basmati | Basmati Type III | India | Europe |
|  | Super Basmati | Pakistan | Europe |

(\*) Whole-genome sequencing data retrieved from <sup>14</sup>

### Sequencing, preprocessing, and mapping

DNA libraries were prepared with Truseq DNA PCR-free protocol and whole-genome sequencing (WGS) was performed using the Illumina NovaSeq 6000 platform (Macrogen, South Korea). Raw paired-end read quality was assessed using FastQC (v0.11.9) ^20^. Due to the high quality of the reads, alongside the absence of adapters, no read trimming was applied for most accessions. Exceptionally, for Bomba and Puntal (reads obtained from ^14^), low-quality nucleotide regions were removed (QS ≥20) using Trimmomatic (v0.39) ^21^

High-quality reads were then mapped to the reference genome Nipponbare 1.0 (IRGSP-1.0, release 52) using bwa-mem v0.7.17 with default parameters ^22^. The resulting SAM files were converted into BAM files (option ‘samtobam’) and coverage/depth statistics were obtained using option ‘depth’ from SAMtools v1.7 ^23^. Additionally, BAM files were sorted (option ‘sort’) and indexed (option ‘index’), and duplicate reads were marked (option ‘dedup’) using SAMtools.

### Variant calling and filtering

Short variant calling was performed using Genome Analysis Toolkit 4 (GATK v4.2.6.1), following the GATK “Guide of good practices for the discovery of germline short variants” ^24^. In detail, quality scores for each base pair were recalibrated using known sites of *O. sativa* variants (https://ftp.ensemblgenomes.ebi.ac.uk/pub/plants/release-60/variation/vcf/oryza_sativa/oryza_sativa.vcf.gz). Then, SNPs and InDels were called for each variety using GATK HaplotypeCaller, and stored in genomic variant calling format (gVCF) files. These 22 files were merged by joint genotyping in a single cohort VCF file using GATK CombineGVCFs and GenotypeGVCF options.

SNPs and InDels records were stored in 2 separate files using GATK SelectVariants for filtering them independently, as recommended in the GATK best practices and as previously reported by Ji *et al*. ^25^. Low-quality SNPs were removed based on: allele depth (QD) < 5, strand bias estimated by Symmetric Odds Ratio test (SOR) > 3, Fisher exact test (FS) > 50, root mean square of the mapping quality of reads across all samples (MQ) < 50, rank sum test for mapping qualities (MQRankSum) < -2.5 and the relative positioning of reference versus alternative alleles within reads (ReadPosRankSum) < -1.0 and > 3.5. Low-quality InDels were removed based on QD < 2.0, FS > 200.0, and ReadPosRankSum < -20.0.

### Estimation of variant effects

A field containing annotations regarding the position (related to annotated genes) of each variant, in addition to their putative effect (HIGH, LOW, MODERATE, MODIFIER) on gene function was added to the files containing each type of variant, using SnpEFF v5.1^26^, using the built-in structure annotation library for *Oryza sativa* (MSU7). A functional enrichment analysis of the identified genes annotated with HIGH impact SNPs was performed using the gprofiler2 R package (‘gostres’ and ‘gostplot’ functions) ^27,28^

### Phylogeny analysis

The phylogenetic tree was generated with the total filtered SNPs in VCF2PopTree ^29^ using default parameters (-output “Newick tree”). Then, tree labels were colored according to the rice commercial category, and exported in .svg format using the Interactive Tree of Life (v6) online tool.

### QTLs and gene enrichment analysis

Start and end positions of Quantitative Trait Locus related to rice-eating quality and seed properties were retrieved from the Rice SNP-Seek database (last accessed on January 31^st^, 2023). Using a custom R script (*snp_in_qtl*.*R*), genes annotated with HIGH impact SNPs were screened to check if their variants occur within the collected QTLs of interest. Then, the resulting table was used for a gene enrichment analysis using the gprofiler2 R package (custom R script *variant_enrichment*.*R*).

## Results

### Genome-wide profiling of commercially valuable varieties from the Mediterranean market

To obtain the genetic polymorphisms present in the genome of the commercially valuable varieties selected in this study, high-quality paired-end reads obtained for each genotype were aligned to the rice reference genome (Nipponbare). For all 20 varieties sequenced within this study, the average percentage of high-quality reads mapped and properly paired in unique positions was 97 %. As a result, on average, over 127 M of reads were unique alignments (Fig. 1a) with a final mapping depth ranging from 46.0x (Maçarico) to 62.6x (Manobi), with a mean of 53.0x coverage. For Bomba and Puntal varieties, about 98.2% of the total reads were mapped and paired and 87% were unique alignments. A final coverage of 30.5x for Bomba and 25.8x for Puntal (Table 2) was obtained. Given the good quality of the paired-end read files and the high coverage after alignment to the reference genome, the final BAM files were prepared for short variants extraction. These files included both those generated in this study and previously published sequencing data. The full array of SNPs and InDels obtained using GATK was filtered, leading to the identification of over 4.8 M high-confidence variants, of which over 3.6 M were SNPs and 1 M were InDels (Table 2). From the sequencing data, out of the 12 rice chromosomes, chromosome 11 had the highest number of variants, contrasting with chromosome 9 (the smallest one). The size of the chromosomes was taken into consideration for the calculation of variant rate and density, which revealed chromosome 5 as the one with the least number of variants per 1 kbp window (density) and a higher rate (Table 2), which describes the mean number of base pairs in which one variant occurs ^25^. Additionally, an analysis of the obtained substitutions, in the case of SNPs, and of length, in the case of InDels, was performed (Fig. 2b,c). It is noticeable that most SNPs corresponded to a substitution of Cytosine to Thymine (C>T) or Guanine to Adenine (G>A) (Fig. 2c). Regarding InDels, the comparison with the reference genome showed a higher predominance of shorter InDels (<5 bp), which was an expected result.

**Table 2.**
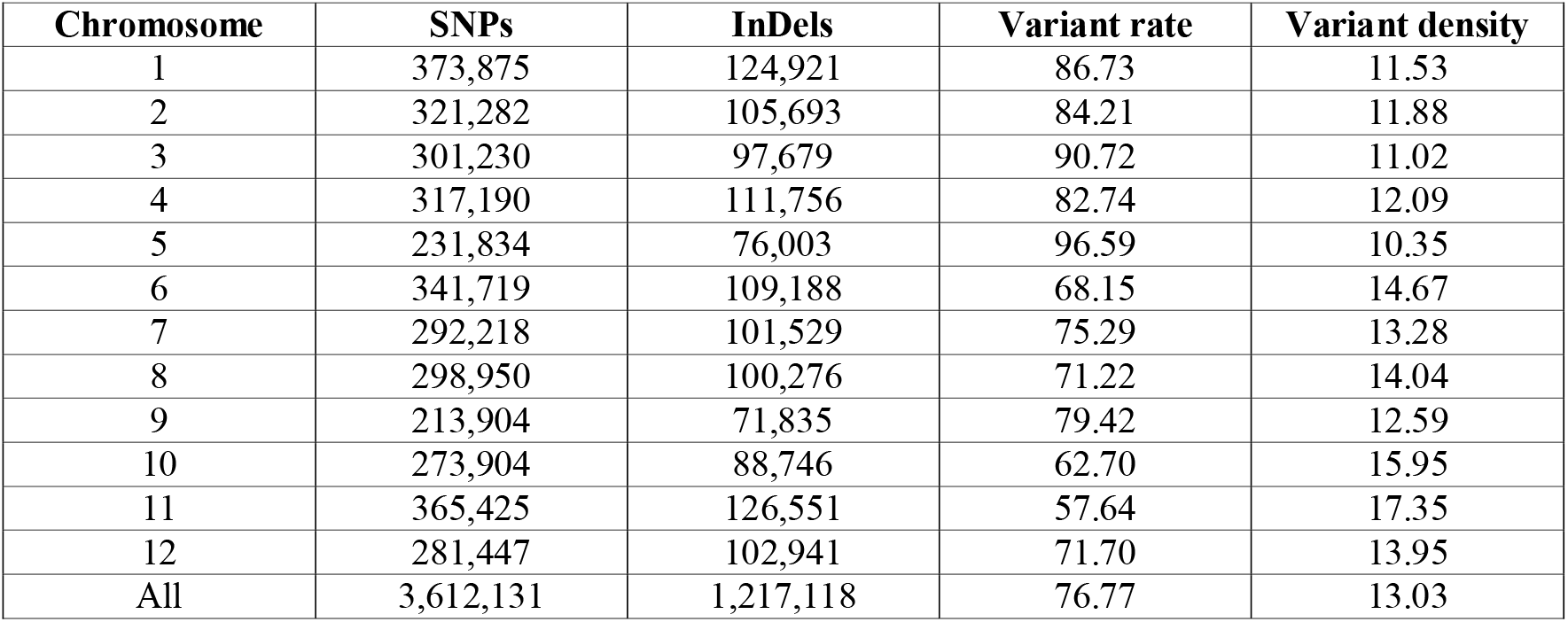
Table with the number of variants (SNPs and InDels), variants’ rate, and density, after filtration with the filters suggested by ^25^. Rate is the mean of base pairs in which one variant occurs and density considers the mean of variants that occur in a window of 1 kbp. All are based on chromosome size (sequencing data).

| Chromosome | SNPs | InDels | Variant rate | Variant density |
| --- | --- | --- | --- | --- |
| 1 | 373,875 | 124,921 | 86.73 | 11.53 |
| 2 | 321,282 | 105,693 | 84.21 | 11.88 |
| 3 | 301,230 | 97,679 | 90.72 | 11.02 |
| 4 | 317,190 | 111,756 | 82.74 | 12.09 |
| 5 | 231,834 | 76,003 | 96.59 | 10.35 |
| 6 | 341,719 | 109,188 | 68.15 | 14.67 |
| 7 | 292,218 | 101,529 | 75.29 | 13.28 |
| 8 | 298,950 | 100,276 | 71.22 | 14.04 |
| 9 | 213,904 | 71,835 | 79.42 | 12.59 |
| 10 | 273,904 | 88,746 | 62.70 | 15.95 |
| 11 | 365,425 | 126,551 | 57.64 | 17.35 |
| 12 | 281,447 | 102,941 | 71.70 | 13.95 |
| All | 3,612,131 | 1,217,118 | 76.77 | 13.03 |

**Fig. 1.**
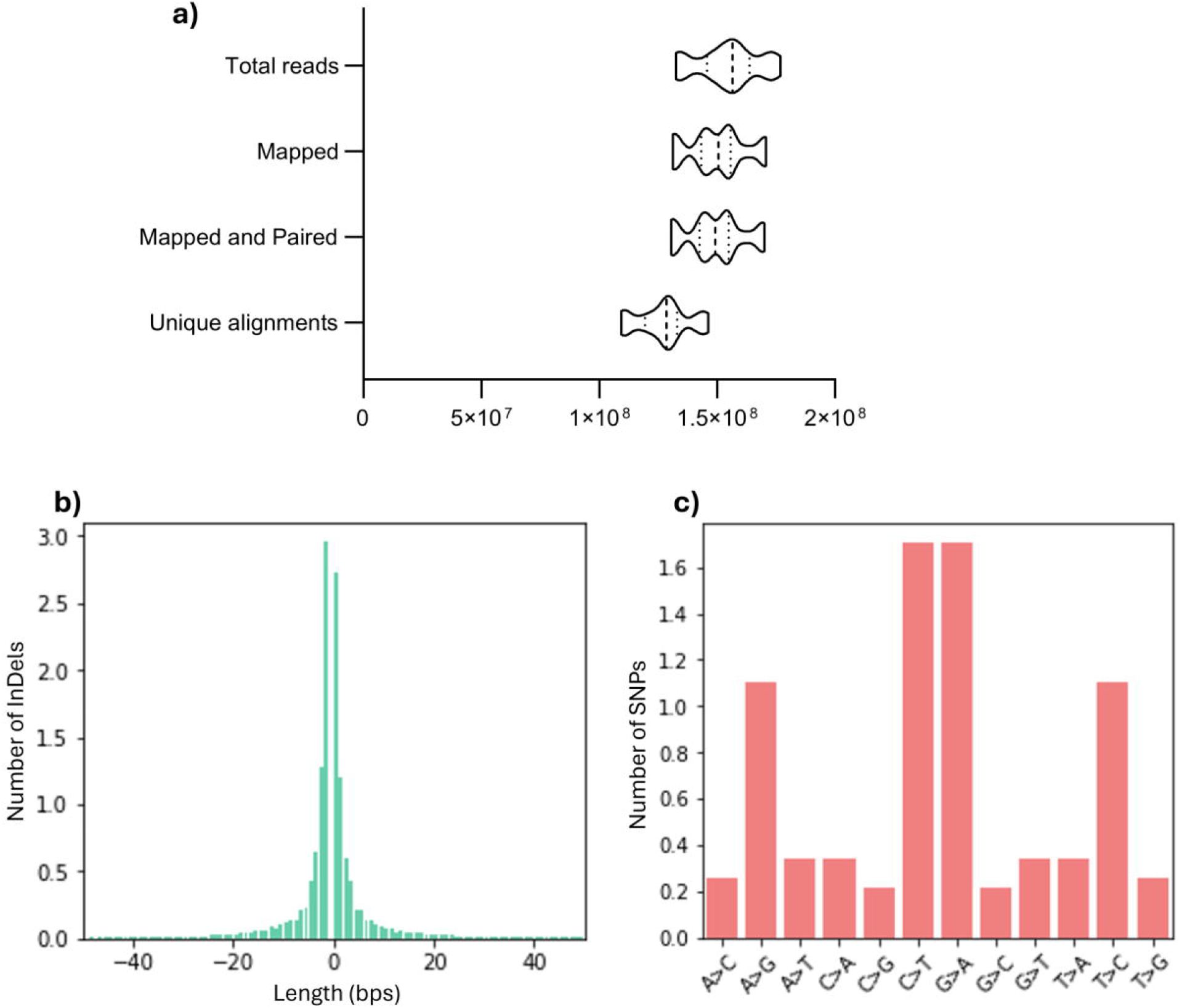
Whole-genome sequencing of 20 varieties and the characteristics of the generated polymorphisms, with the addition of two Spanish genotypes. **a)** Number of reads obtained from the whole-genome sequencing of 20 rice varieties. The violin plot represents the distribution of total reads, mapped reads, mapped and paired, and unique alignments obtained from the alignment with the rice reference genome. **b)** Length of the InDels identified in 22 rice varieties, in comparison with Nipponbare. Negative numbers correspond to deletions and positive to insertions of a number of nucleotides. The number of InDels with each size is on a logarithmic scale. **c)** Substitutions detected on SNPs data. A- Adenine, C- Cytosine, G- Guanine, T- Thymine. The number of SNPs is on a logarithmic scale

**Fig. 2.**
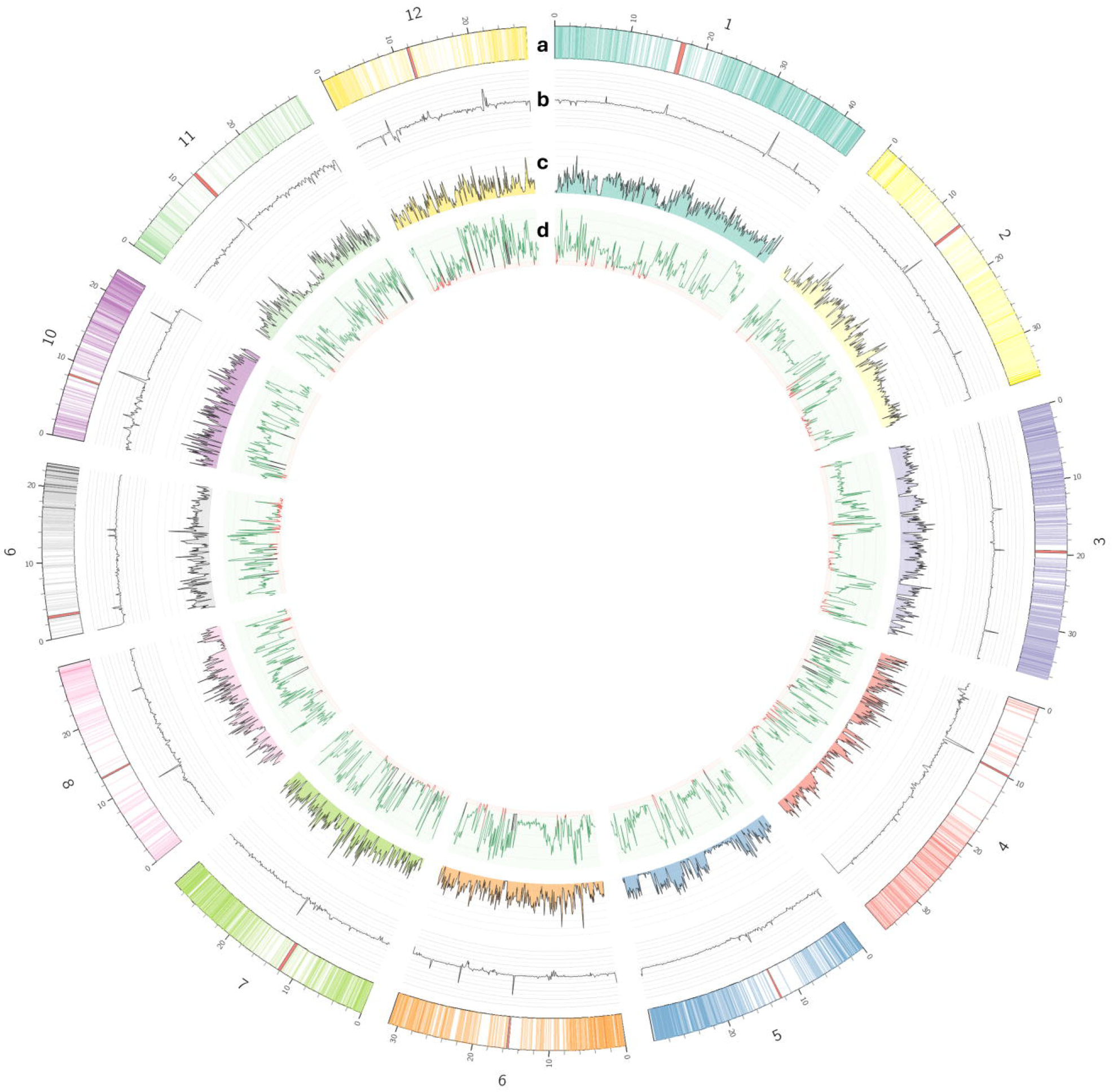
SNPs analysis at whole-genome level. Circos plot represents (from the outer to the inner ring): **a)** Chromosome size, centromere position (red band), and coding genes density (by intensity of each color); **b)** coverage of reads per chromosome; **c)** SNPs density (number of SNPs in 100 Kb windows); **d)** Tajima’s D mean in each 100 Kb window, with positive values in green and negative values in red

### Variant Information

A comprehensive analysis of the SNPs along the twelve chromosomes in rice highlighted a somewhat uneven distribution (Fig. 2). It should be noted that no correlation between SNPs occurrence and sequencing coverage was found, indicating a reduced bias from underrepresented regions (Fig. 2b,c). Although some peaks of coverage occur in parts of the genome with lower SNP detection, this is not observed in the majority of peaks or the chromosomes, for instance when comparing chromosome 4 to chromosome 10 (Fig. 2b,c). Also, when examining the gene density, there seems to be no correlation with SNP distribution. Although some of the less dense regions of each chromosome correspond to the centromere, those regions have less occurrence of SNPs in most chromosomes. Tajima’s d was calculated to further reveal regions of the genome where the observed variation is conserved. A few chromosome regions (end of chromosome 1 and chromosome 10) are highlighted by having high Tajima’s *d* positive peaks (Fig. 2d), suggesting an event of balancing selection.

### SNP density across each chromosome reveals conserved polymorphisms between varieties

The whole-genome SNP-based phylogenetic tree highlights two main groups. The first cluster groups together both Basmati varieties (Super-Basmati, Basmati Type III) alongside Giza181 and Maçarico. The second cluster groups all other varieties, with subsequent divisions highlighting Long B varieties (CL-28, Puntal) grouping, as well as the medium grain varieties (in green), JSendra, and Manobi. Based on SNP data alone, and lacking the pedigree information for these varieties, we suggest that Giza 181 and Maçarico are more closely related to Basmati varieties than the others assessed in the study, despite belonging to separate categories defined from phenotype observation.

When observing SNP density along each chromosome for all varieties, we found that Maçarico, Giza 181, Basmati Type III, and Super Basmati clustered together with a tendency to accumulate a higher number of SNPs (Fig. 3). This trend was conserved in most chromosomes (Fig. 4), with the exception of Chr9. This particular group of 4 varieties is more evident in 9 of the total chromosomes (Chr1 - Chr5, Chr8, and Chr10 - Chr12), with 3 of them (Maçarico, Giza 181, and Basmati Type III) always grouping within the same cluster in all chromosomes. The clustering of the remaining varieties based on SNP density per chromosome was less conserved, with the overall relatedness being chromosome-dependent.

**Fig. 3.**
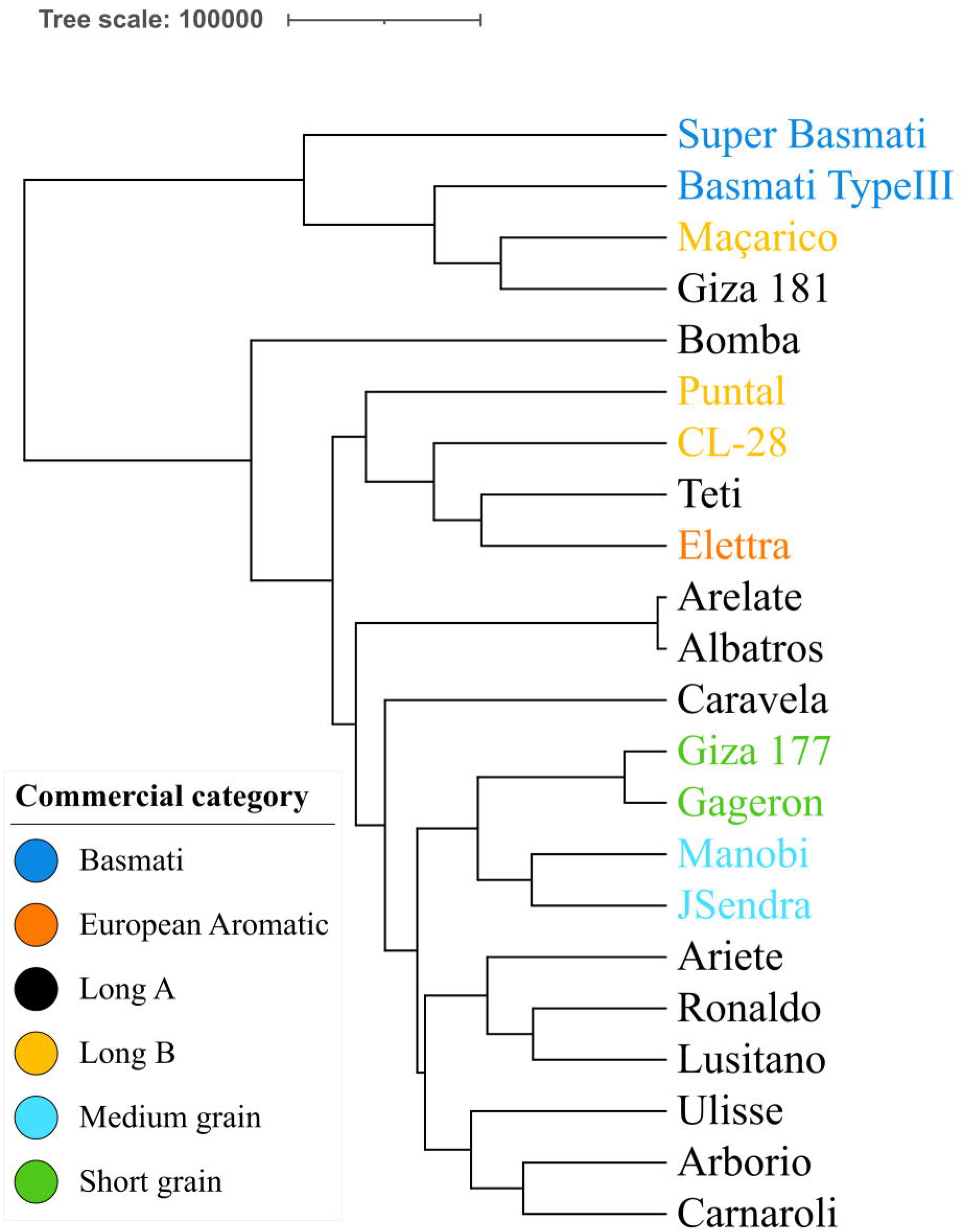
SNP-based phylogeny of the 22 varieties. Color labelling is based on defined rice commercial categories

**Fig. 4.**
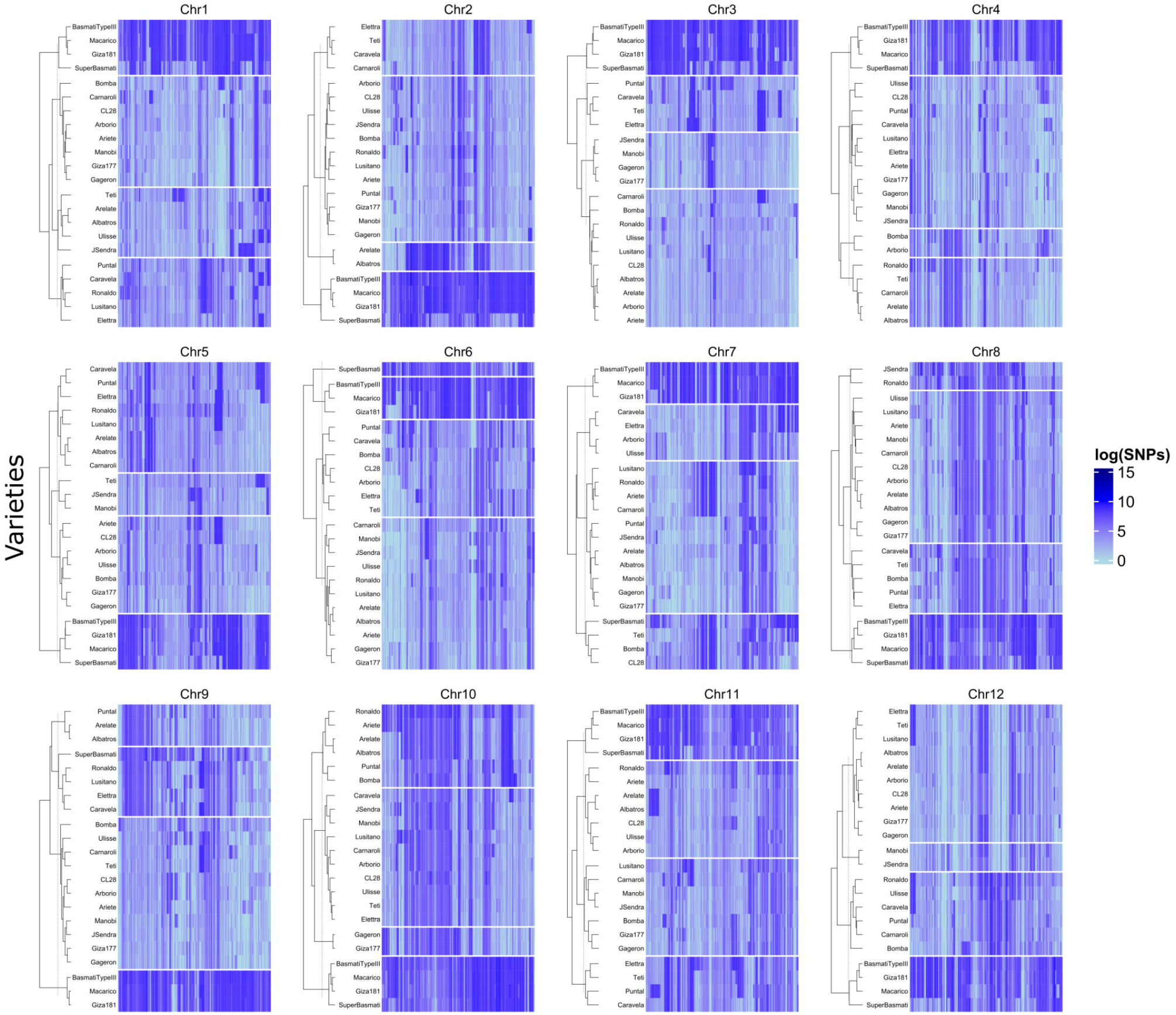
SNP distribution in the 12 rice chromosomes for all 22 varieties, with hierarchical clustering (k=4) highlighting groups with similar distribution patterns

### Global and variety-specific genes in QTLs of interest

The list of collected SNPs was annotated according to their putative impact on the genomic sequence. To assess the functional impact of the detected diversity, we further identified high-impact variants, predicted to disrupt a start codon or introduce a stop codon within the coding sequence of genes in all varieties targeted in this study. A total of 1003 high-impact SNPs was identified (Online Resource 1) associated with 912 genes. This list of genes was subsequently filtered for genes within previously identified QTL regions related to eating quality traits and/or seed properties, resulting in 911 genes. Table 3 highlights a group of genes with particular interest due to their previously described roles in regulating seed traits, and specific variants within their coding sequences which may actively impact these traits.

**Table 3.**
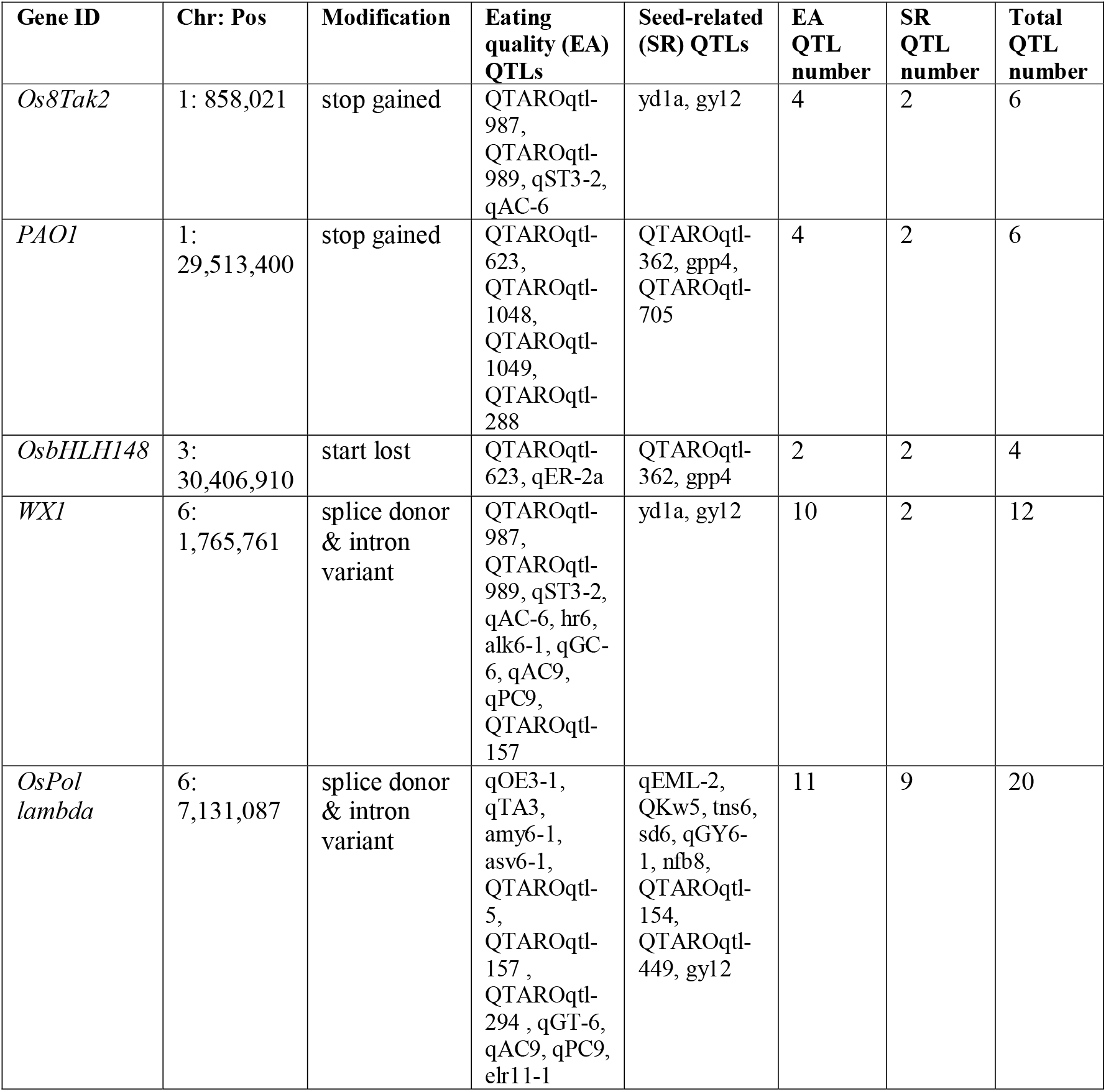
Genes of interest with HIGH impact SNPs (annotated with snpEff^26^) within eating quality and seed-related QTLs. Full table available at Supplementary Material 1.

### Enrichment analysis

The set of genes with one or more HIGH-impact SNPs within their coding sequence was enriched in molecular functions related to ADP and nucleotide binding, and biological processes related to defense response, namely against other organisms. Some examples include the *Os8Tak2* gene, related to disease resistance, the DNA damage repair *OsPol lambda* gene, and the *OsbHLH148* gene, which is involved in abiotic stress tolerance (Fig. 5).

**Fig. 5.**
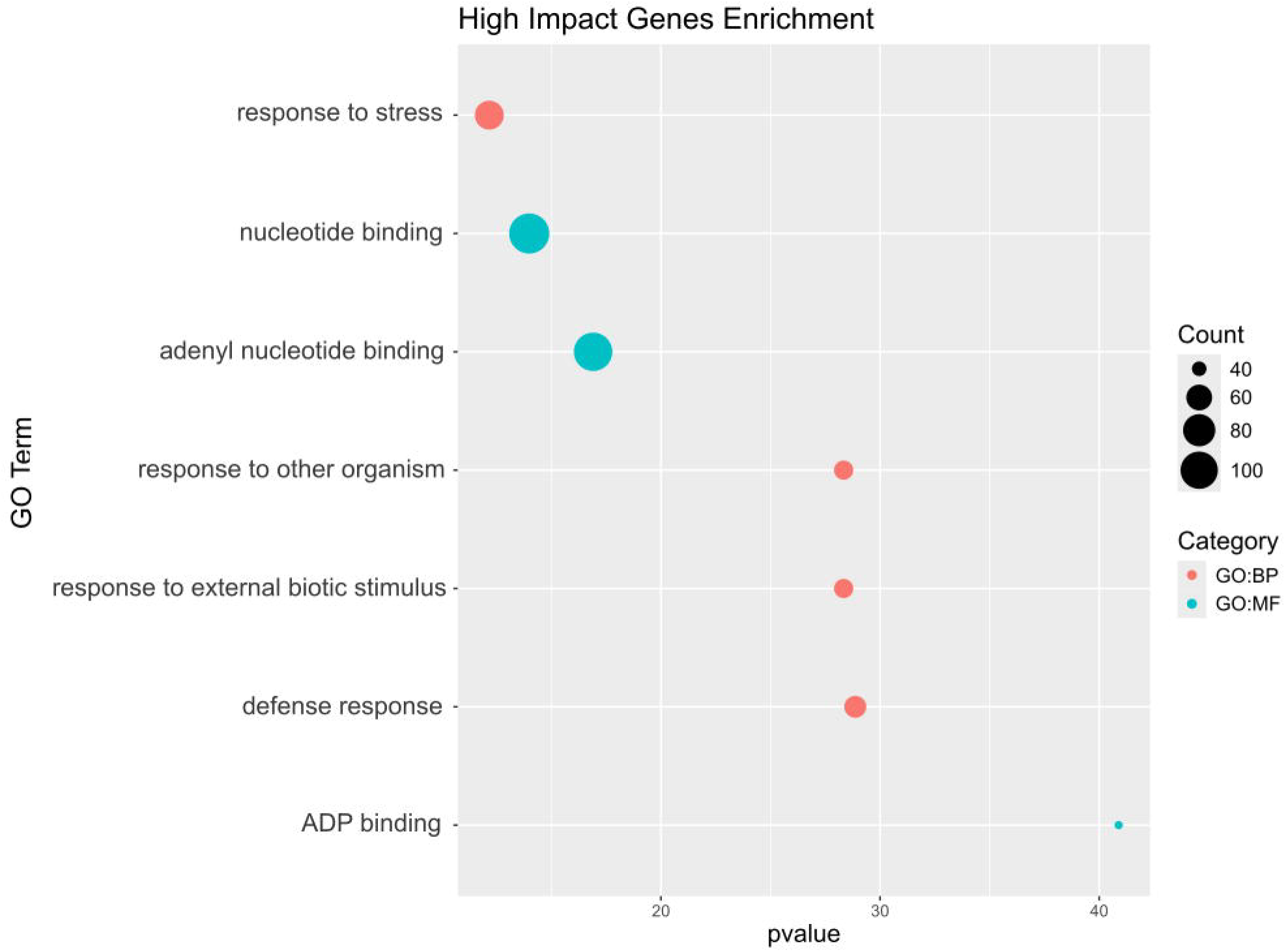
Functional enrichment analysis of genes with HIGH impact SNPs.

## Discussion

In this study, we sequenced 20 and obtained the global polymorphisms for 22 rice varieties, produced and or circulating in the Mediterranean region. This list was selected based on factors such as market value and the rice potential for breeding applications.

The SNP distribution at the chromosome level highlights that the 22 targeted varieties form clusters that are mainly consistent across the genome. Phylogenetic inference from SNP density and variability enabled the prediction of genetic relatedness among these varieties. This analysis divided the group into two main clusters, one with both Basmati varieties, plus Maçarico (Long A) and Giza181 (Long B). This shows that the high-quality polymorphisms dataset generated in this study has no direct link with their respective commercial category, which is mainly based on grain-related phenotypical traits. We hypothesize that this direct correlation may not be reached without the full context of varietal pedigree of the varieties (undisclosed by seed providers), and putative epigenetic impact.

Interestingly, out of the 912 genes annotated with one or more HIGH impact SNPs, 911 of them located within QTLs of interest related to eating quality and seed-related properties. These polymorphisms were identified in genes such as Waxy, *OsPol lambda*, and *PAO1*, and are present in at least one of the varieties in this study. The well-established *Waxy (Wx)* “granule-bound starch synthase 1” gene is responsible for amylose synthesis, a key determinant of rice cooking and processing qualities ^16^. This gene is located in chromosome 6 and is genetically linked to Eating Quality QTLs (QTAROqtl-157, QTAROqtl-987, QTAROqtl-989, qST3-2, qAC-6, hr6, alk6-1, qAC-6, qGC6, ac6, qAC9, qPC9) and seed related QTLs (yd1a, gy12). In the endosperm, the expression of this gene is highly impacted depending on different alleles within the Wx locus, which leads to differential amylose content. This allele forms relate to a transversion (T-G) located in pos-1,765,761bp that impacts the splicing of the Wx transcript, leading to changes to amylose content. The presence of the G allele correlates with higher amylose content and the T allele is linked to lower amylose content ^16^. Interestingly, we confirmed the presence of this SNP in both Basmati varieties targeted in this study, in addition to Maçarico, Carnaroli, CL28 (heterozygous) and the Spanish varieties Bomba and Puntal (homozygous for the alternate allele). This matches the phenotypic trait of Basmati varieties, which have lower starch but higher amylose content within their grain ^30^.

It was previously shown that, when over-expressed, the *OsbHLH148* gene confers drought tolerance in rice ^31^. We have identified a high-impact SNP within the coding region of this gene (pos-chr3: 30,406,910bp) in the Albatros, Arelate, Ariete, Lusitano, and Ronaldo varieties. This SNP is annotated as disrupting the single transcript start codon, which may lead to the gene’s defective transcription and compromise its role in drought response. Another gene of interest was the *OsPol lambda* (*Os06t0237200-01*) “DNA polymerase lambda” gene, which belongs to the only X family of DNA polymerase in rice involved in DNA damage repair. It is described as up-regulated in response to abiotic stress (drought, salt), and correlating with the stress intensity ^32^. In four of the varieties under study (Albatros, Arelate, Ariete, and Ronaldo), we found a homozygous SNP corresponding to a transition A→G, which was predicted to interfere with the transcript splicing process. This subset of varieties also showed the SNP variant described above for *OsbHLH148*. Collectively, and since drought tolerance represents a trait of interest in the context of climate change, more particularly in the Mediterranean region, we suggest that both genes can be further studied in the high-value varieties produced in this region to better understand specific stress responses during production.

The *Os8Tak2* “receptor-like kinase 20” gene has been previously shown to negatively regulate rice resistance to bacterial blight ^33^ For all varieties except Albatros, Arelate, Caravela, Gageron, Giza181, JSendra, Maçarico, and Manobi, we identified a high-impact SNP annotated as introducing an early stop codon in this gene coding region. The functional enrichment analysis revealed that the group of 911 genes is significantly enriched in processes like stress response and response to other organisms. We suggest that the *Os8Tak2* gene, along with the *OsbHLH148* and *OsPol lambda* genes described above, serve as key examples within this group, potentially playing important roles in the stress response and adaptation of these varieties to the biotic and abiotic factors commonly found in the Mediterranean region.

Rice contains seven genes that encode polyamine oxidases (PAOs), named Os*PAO1* to OsPAO7 according to chromosome and gene ID number. *PAO1*, located in chromosome 1, has a known function in the back-conversion of spermine and thermospermine into spermidine and is described as responding to cytokinin levels in rice ^34,35^. In *Arabidopsis*, the same role is described for its ortholog *AtPAO5*, in addition to promoting seed germination ^36,37^. Within the coding sequence of the *PAO1* gene, we identified a high-impact SNP (chr1:29,513,400) for both Basmati varieties (heterozygous alleles) and for Maçarico, Giza181, Bomba, and Puntal (homozygous for the alternate allele). This SNP is annotated as introducing a stop codon within the gene coding sequence, likely compromising the *PAO1* transcriptional process. In the targeted varieties, *PAO1* occurs in a region with a positive tajimaD value (‘>2’), therefore we hypothesize that it may be undergoing a process of balancing selection and acting as a main discriminator between the two main observed phylogenetic clusters.

In conclusion, we believe that the open availability of the data generated in this study will be useful for researchers and breeders to deepen their knowledge regarding these high-value cultivars, predicted to increase in relevance given the production and consumption trends observed in the Mediterranean region. It may also support further studies focusing on their adaptation potential to the changing Mediterranean climate.

## Acknowledgments

The authors thank the TRACE-RICE project for supplying the biological material, and Biodata.pt for guidance on data management.

## Data availability

Whole-genome sequencing data generated and analyzed during the current study are available in the ENA repository under the **PRJEB64146** accession code. The raw variants data generated and analyzed during the current study are available in the EVA repository under the **PRJEB83571** accession code. Custom scripts used for data analysis during the current study are available on GitHub (https://github.com/hmrodrigues99/TRACE-RICE). Supplementary data and MIAPPE information for sequenced varieties can be found in Dataverse (https://dmportal.biodata.pt/dataverse/gvtritqb).

